# Polyacrylamide-based Hydrogel Coatings Improve Biocompatibility of Implanted Pump Devices

**DOI:** 10.1101/2022.12.13.520347

**Authors:** Doreen Chan, Caitlin L. Maikawa, Andrea d’Aquino, Shyam S. Raghavan, Megan L. Troxell, Eric A. Appel

## Abstract

The introduction of transcutaneous and subcutaneous implants and devices into the human body instigates fouling and foreign body responses that limit their functional lifetimes. Polymer coatings are a promising solution to improve the biocompatibility of such implants, with potential to enhance *in vivo* device performance and prolong device lifetime. In this work we sought to develop novel materials that can be used as coatings on devices. We first synthesized a library of polyacrylamide-based copolymer hydrogels and implanted them into the subcutaneous space of mice to evaluate their biocompatibility. We sought to discover copolymer hydrogel formulations eliciting less fibrous capsule formation and lower overall inflammation than gold standard materials such as PEG and polyzwitterions. The top performing copolymer hydrogel materials were then shown to improve the biocompatibility of a commonly used medical material, PDMS. Lastly, we applied the leading hydrogel coating to the catheter tips of insulin pumps. In a rat model of insulin deficient diabetes, the functional lifetime of hydrogel-coated devices was extended over uncoated pumps. These polyacrylamide-based copolymer hydrogel coatings have the potential to improve device function and lifetime, thereby reducing the burden of disease management for people regularly using implanted devices, such as those with diabetes.

## 1. Introduction

Remarkable progress has been made over the past few decades on long-term implantable devices such as cardiac pacemakers, hernia meshes, and dental and heart implants, which has improved patient health across numerous disease indications. The worldwide medical device industry is estimated to be $150 billion^1^ and uses materials fabricated with metals (e.g., titanium, nitinol, and stainless steel^2^) or soft materials (e.g., polytetrafluoroethylene, silicones, poly(ethylene terephthalate), among others^3^). Often these devices are designed to integrate with surrounding tissue, whereby a gradual process of interfacing with the body is typically mediated by the immune response. Unfortunately, for temporary implantable devices and biosensors such as glucose monitors, insulin pumps, catheters, and neural probes, these immune responses are typically undesirable as they inhibit device performance, shorten device lifetime, cause scarring in the local tissues, and result in a requirement for frequent device replacement^1, 4^.

Upon implantation, materials and devices form a complex interface with tissues^5^ and their surface is instantaneously coated with proteins such as fibrinogen, IgG, fibronectin, and von Willebrand factor^5^. The adsorbed proteins undergo conformational changes and the protein covered surface is activated — behaving as a receptor and initiator site — and initiating a foreign body response (FBR). The FBR is an acute inflammatory response that dictates how surrounding immune cells respond, including attracting other cells or masking the material^6^. This process is dynamic as cells, especially macrophages, are recruited to the implanted materials and devices and secrete more proteins and respond directly to the foreign material. Over time, fibroblasts secrete collagen fibrils on the surface of these implanted materials and devices that results in the formation of a fibrous capsule that is avascular, impermeable to cells, hinders metabolite transport, and serves to isolate the material from the body. This fibrotic to implanted materials ultimately compromises long-term function of implantable materials and devices^7, 8^.

Devices used for effective management of type I diabetes mellitus (T1D) are particularly affected by the immune system and resulting foreign body response. People with diabetes rely on continuous glucose monitors (CGMs) to evaluate glucose levels in real time, as well as insulin pumps to provide basal and bolus infusions of insulin to manage blood glucose. Both of these devices utilize a catheter tip that is implanted in the subcutaneous space of the patient’s abdomen. While insulin pumps face many technical challenges, such as battery life and viability of insulin on board, the most burdensome challenge for their effective use is occlusion of the infusion set catheter tips on account of the FBR^9^. Indeed, aggregation of administered insulin at the catheter tip encourages and exacerbates the FBR, resulting in occlusion and flow irregularities. For many insulin drug products, infusion sets are recommended to be removed and replaced at a new site in the patient’s abdominal subcutaneous tissue at a minimum of every 3 days. This highly burdensome process leads to noncompliance among those with diabetes, and people who use infusion sets for longer than the recommended period can experience irritation at the insertion site and inconsistent insulin delivery resulting in poor disease management^10^. Thus, there is a strong motivation to improve the biocompatibility of the inserted catheter tips to mitigate occlusion and extend the lifetime of these crucial devices.

To address the shortcomings of current infusion set catheters, significant efforts towards surface and bulk modifications have been made to these implanted devices^5, 11^. In particular, antifouling polymeric brush and hydrogel coatings are an attractive option for surface modification, as they serve to mediate the interface between implanted materials and the body. These modifications can be designed to prevent adsorption of proteins and preclude infiltration of immune cells to reduce the severity of the FBR^12,13^. Notably, hydrogels comprising zwitterionic monomers such as poly(2-methacryloyloxyethyl phosphorylcholine) (PMPC) have been shown to exhibit a high degree of biocompatibility^14, 15^, and PMPC-based polymer brushes have demonstrated an ability to improve performance and reduce noise on continuous glucose monitors^16^. Similarly, poly(ethylene glycol) (PEG) is often regarded as the gold standard polymer for antifouling coatings^18–21^. Unfortunately, due to rising concerns of PEGs immunogenicity and recent research demonstrating that both PEG and PMPC exhibit poor long-term stability, there is growing interest in the discovery and development of new classes of antifouling materials. Recently, polyacrylamide-based copolymer hydrogel coatings have been reported which exhibit excellent anti-biofouling properties and stability^17^. In this work, we investigate a series of antifouling polyacrylamide-based copolymer hydrogel materials to identify coating materials that reduce the FBR to implanted materials, including catheters on insulin pumps. We show that select hydrogel formulations mitigate the FBR and improve the functional lifetime of implanted insulin pump devices.

## 2. Results

### 2.1. Selection and Synthesis of Polyacrylamide Hydrogels

While the link between fouling and biocompatibility is still poorly understood^18^, it has been generally agreed upon that protein adhesion is the first step in fouling. This protein fouling is followed by infiltration of neutrophils (acute inflammation), recruitment of macrophages and monocytes (chronic activation), and cell fusion to form foreign body cells. This process is enhanced by secretion of soluble factors and collagen production by fibroblasts (**Fig. 1**). From a recent study evaluating a large library of polyacrylamide-based copolymer hydrogels for their ability to prevent founding and platelet adhesion^22^, we selected the top 15 antifouling formulations (**Table 1, Fig. S1**) to compare their biocompatibility following subcutaneous implantation in mice against PEG and zwitterionic PMPC hydrogel formulations (**Fig. 2a**). Each of these hydrogels can be prepared by photo-polymerization in a facile manner to yield a library of hydrogel discs with mechanical properties which can be tuned to be suitable for interfacing with the body. While we primarily investigated the top 15 anti-biofouling formulations from the previously reported screen of platelet fouling^22^, a subset of randomly selected polyacrylamide-based copolymer hydrogels exhibiting varying levels of antifouling and platelet adhesion behavior were also synthesized and evaluated (**Fig. S2**).

**Figure 1.**
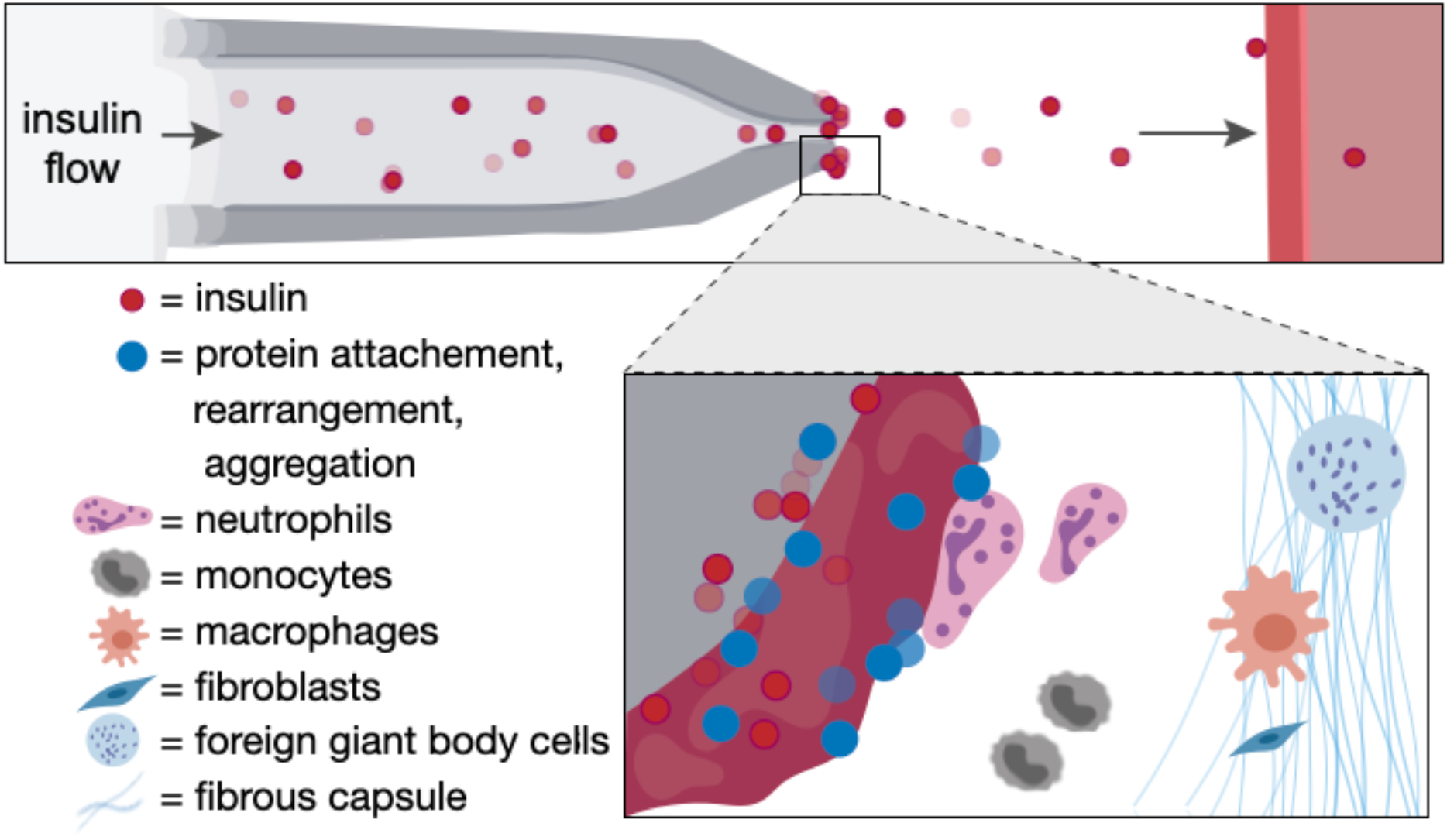
Schematic of the foreign body response to an insulin infusion set. Protein aggregation and innate immune cell activation at the catheter tip of subcutaneously implanted pumps for continuous insulin infusion leads to a cascading foreign body responses that result in device failure, requiring changing of the insulin infusion set.

**Table 1.**
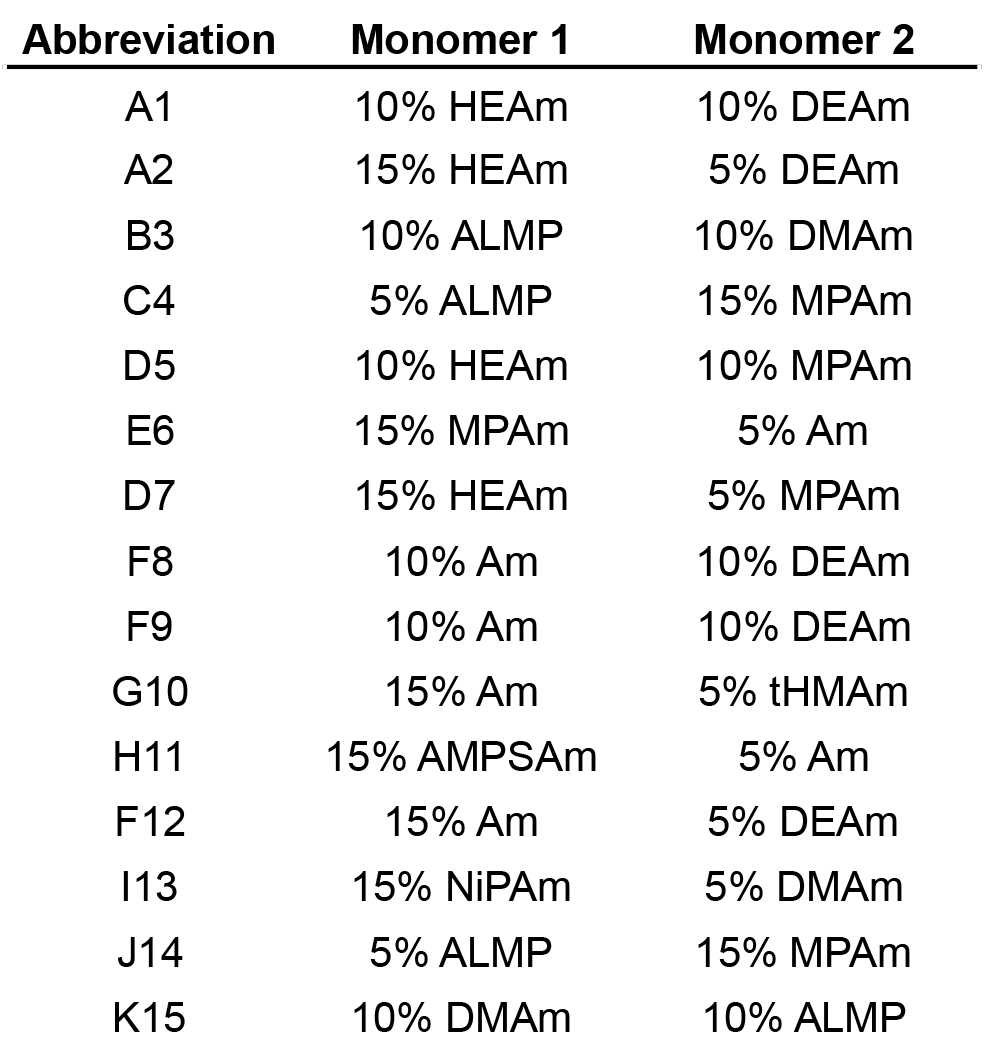
Table of combinatorial polyacrylamide copolymer hydrogel formulations evaluated. Binary mixtures of acrylamide monomers were used to prepare hydrogels (20 wt% solids).

**Figure 2.**
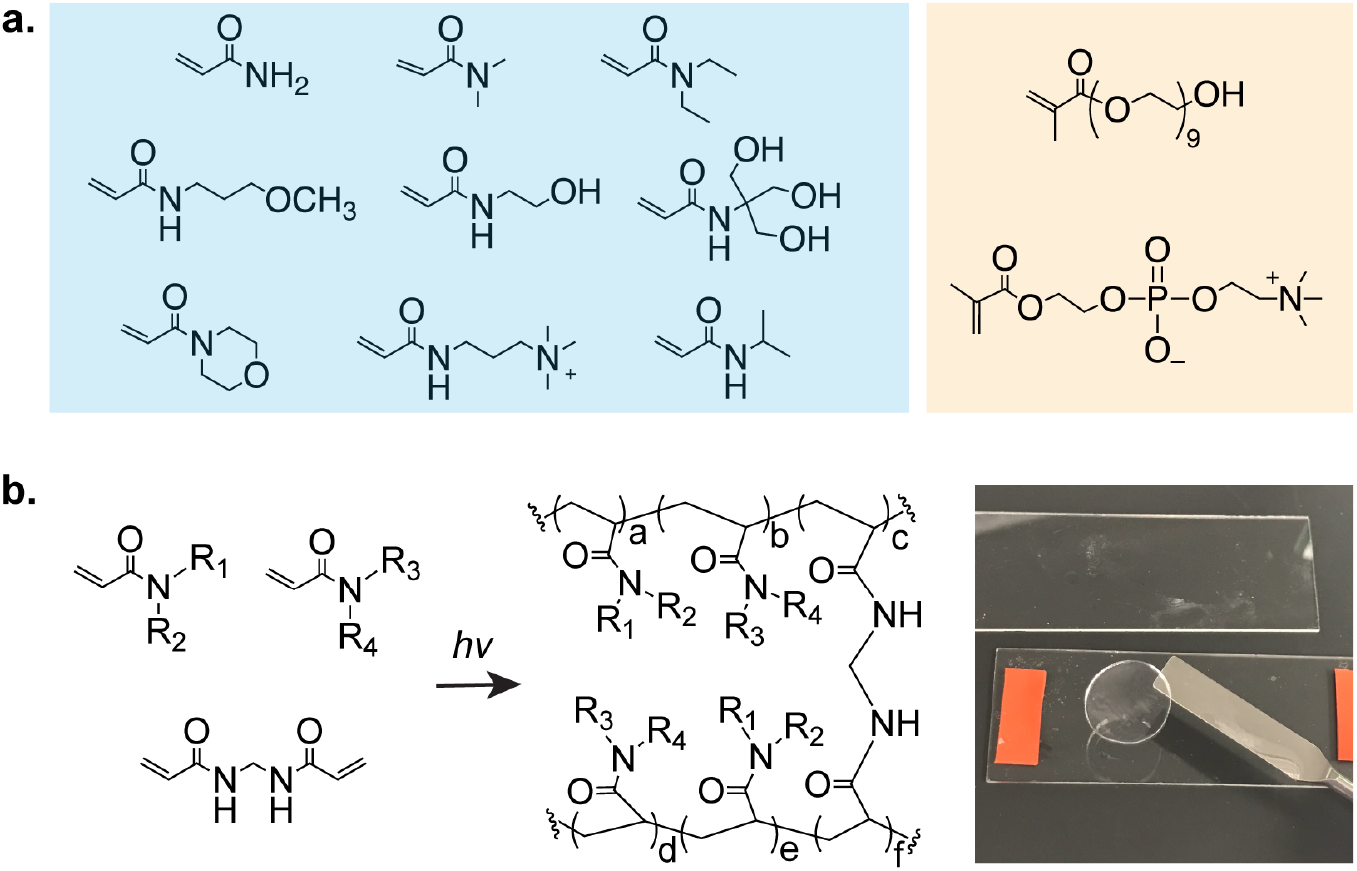
Polyacrylamide copolymer hydrogels to combat the foreign body response. **a**. (left to right) acrylamide (Am); N,N-dimethylacrylamide (DMAm); N,N-diethylacrylamide (DEAm); N-(3-methoxypropyl)acrylamide (MPAm); N-(2-hydroxyethyl)acrylamide (HEAm); N-[tris(hydroxymethyl)methyl]acrylamide (tHMAm); 4-acryloylmorpholine (ALMP); N-[3-(dimethylamino)propyl]methlacrylamide; 2-acrylamido-2-methyl-propane sulfonic acid (AMPSAm); N-isopropylacrylamide (NiPAm). Poly(ethylene glycol) methacrylate and PMPC as controls. **b**. Polymerization of acrylamide monomers are synthesized in a high-throughput manner.

### 2.2. Biocompatibility of Polyacrylamide Hydrogels

As the FBR severely hinders *in vivo* performance of implanted devices, we first sought to evaluate the FBR to each member of our hydrogel library and gold standard controls. To screen these materials, we implanted two hydrogel discs in the subcutaneous tissue in the dorsal region of a mouse. After 28 days, no mice showed signs of discomfort or distress, and implanted hydrogels and surrounding tissues were retrieved for histological analysis (**Fig. 3a**). One measure of the FBR is quantification of the thickness of the collagenous fibrous capsules surrounding implanted materials. After 28 days, excised tissues were processed and stained with Masson’s Trichrome (MT) for evaluation (**Fig. 3b**). With MT staining, collagenous fibrosis stains blue, edema and loose fibrosis stains pale blue, and more organized, dense fibrosis stains darker shades. In contrast, cells generally counterstain red, including inflammatory cells and macrophages that are part of the pseudosynovial layer^19^. Capsule thickness quantification highlighted that a copolymer hydrogel formulation comprising 10 wt% HEAm and 10 wt% MPAm (50:50 ratio of HEAm:MPAm) elicited the thinnest capsule layer (28 ± 11 μm) while gold-standard polymers including PEG (51 ± 16 μm) and PMPC (72 ± 41 μm) elicited much more severe capsule formation (**Fig. 3c**).

**Figure 3.**
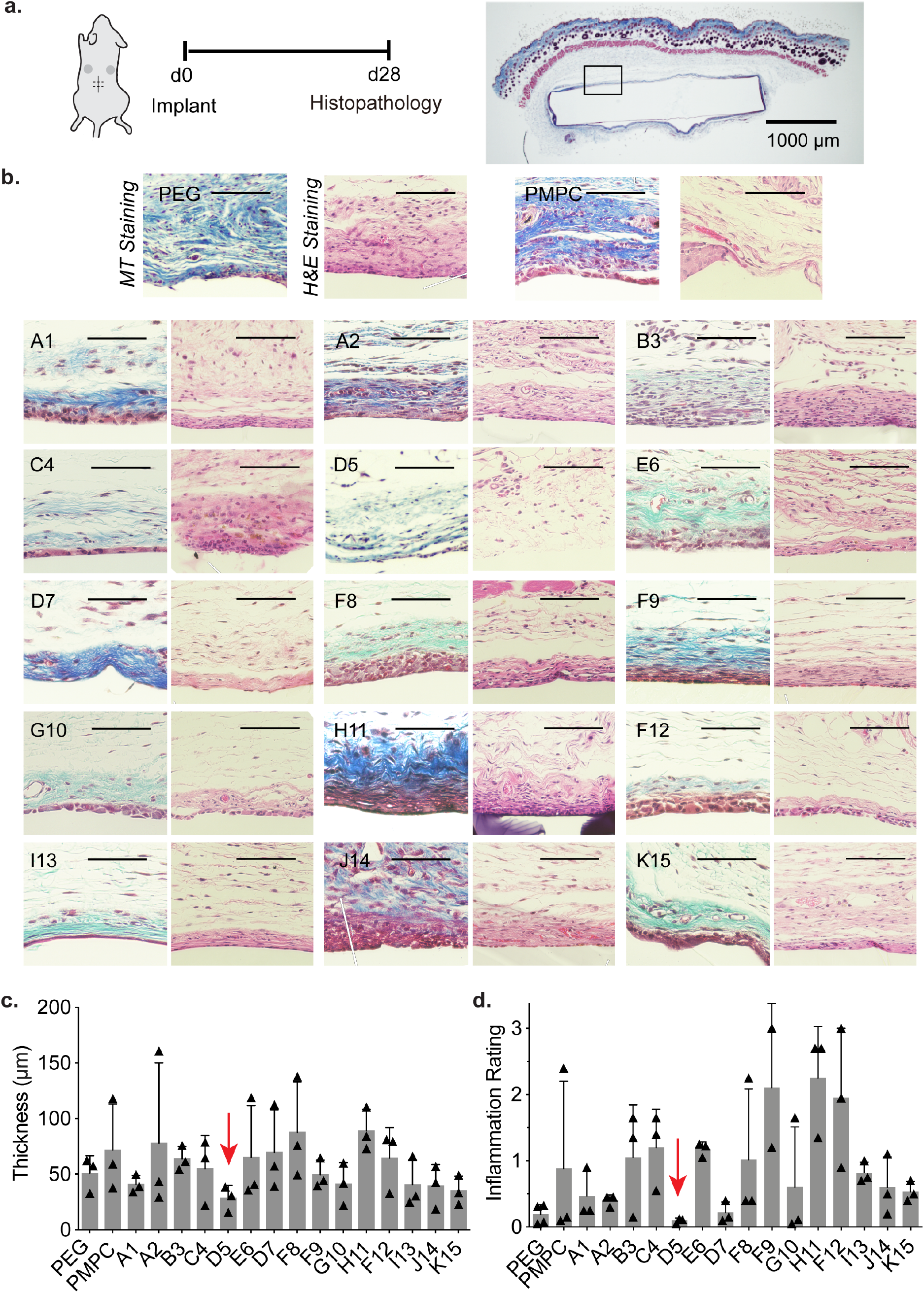
Screening of copolymer hydrogels to select top immune-resistant materials for further evaluation. **a**. Polyacrylamide copolymer hydrogels were implanted in the subcutaneous space of mice for 28 days, upon which the tissues and gels were removed to examine foreign body response. **b**. Representative sections of tissue and subsequent MT and H&E Staining. Scale bar represents 100 μm unless otherwise specified. **c**. Fibrous capsule thickness of n = 3 samples (mean ± s.d.) where each mean was determined from the median of 10 thicknesses per image. **d**. Inflammation score determined by pathologists blinded to the test materials to characterize extent of inflammation.

As fibrous capsule formation provides only one aspect of the FBR, we then investigated what cell types were recruited to the implants as well as the intensity of inflammation near the implants. Excised tissues were then stained with hematoxylin and eosin (H&E), where proteins (e.g., extracellular matrix deposition) stains pink, while nucleic acids (particularly cell nuclei) stains purple (**Fig. 3b**). In all formulations, fibrous capsules were found to be lined by multi-layered collections of histiocytes, forming a pseudo-synovium around the implanted materials. The immune cells surrounding both PEG and PMPC hydrogels were predominantly macrophages. In copolymer hydrogel formulations such as J14, sparse chronic inflammation including plasma cells, lymphocytes and macrophages were present in the stroma deep to the capsule^20^. The presence of neutrophils indicates acute inflammation and was observed to correlate with more severe inflammatory infiltrates, admixed with macrophages and lymphocytes. For hydrogel sample H11, for example, cell presence was dominated by macrophages and neutrophils. In contrast, copolymer hydrogel formulation D5 exhibited few chronic inflammatory cells in stroma, and rare plasma cells and macrophages. A semi-quantitative metric was used to assess the degree of inflammation (neutrophils, lymphocytes, plasma cells, monocytes, giant cells, not including the pseudosynovium) around the hydrogels. For this evaluation, the intensity of inflammation was assigned a score of 0 (none), 1 (mild), 2 (moderate), 3 severe inflammation, which was multiplied by the estimated percent of capsule circumference involved by inflammation. Blinded pathologists scored hydrogel formulation D5 with the lowest inflammation score of all of the materials evaluated: 0.10 ± 0.04 (**Fig. 3d**), In contrast, gold standard materials PEG (0.19 ± 0.15) and PMPC (0.8 ± 1.1) exhibited higher and more variable inflammatory responses. These results indicate that the leading polyacrylamide copolymer hydrogel coating D5 exhibited notably improved biocompatibility over gold standard materials PEG and PMPC.

### 2.3. Application of Hydrogel Coating to PDMS Surface

In the body, proteins readily attach to the hydrophobic surface of polydimethylsiloxane (PDMS), a widely used implant material found in catheters, gastric bands, and breast implants^21, 22^. As a proof-of-concept for the applicability of our polyacrylamide-based copolymer hydrogels as coatings on implanted materials, we applied this hydrogel formulation to a PDMS substrate. Despite prevalent use as a biomaterial historically, these silicone-based materials have been subject of renewed controversy^23–26^. Here, a thin layer of prepolymer solution of our top performing formulation, D5, was applied by spin-coating on PDMS discs and photo-crosslinked to form a robust 46 μm hydrogel coating, as shown by SEM (**Fig. 4a**). Considerably less platelet adhered was observed on the surface of the D5-coated PDMS discs than on bare discs when subjected to platelet-rich plasma (**Fig. S4**). We then implanted hydrogel-coated and uncoated (bare) PDMS samples into the subcutaneous of mice. After 28 days, the implanted materials and surrounding tissues were harvested for histological analysis. MT and H&E staining was performed on samples from these implants (**Fig. 4b**), demonstrating that the D5 hydrogel coating mitigated fibrous capsule formation and reduced the pathological inflammation score for these materials (**Fig. 4c**).

**Figure 4.**
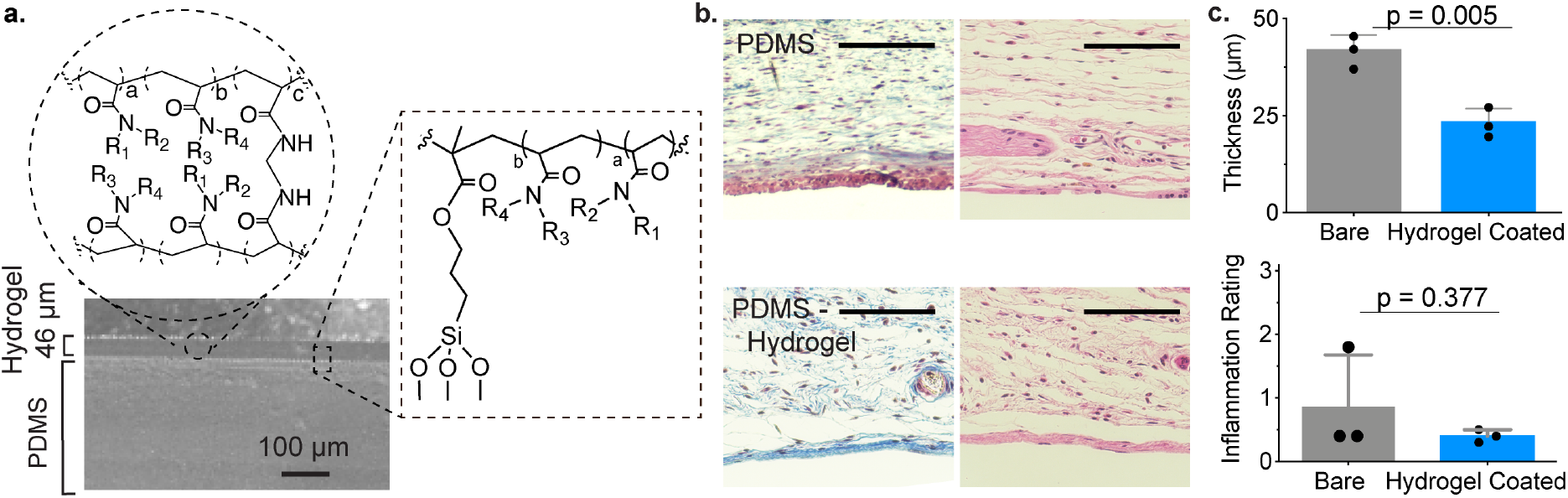
Demonstration of leading hydrogel coating on PDMS surfaces. **a**. Homogenous coating of D5 copolymer hydrogels to PDMS visualized by SEM. **b**. Representative images of MT (left) and H&E (right) staining of explanted hydrogels and adjacent tissues. Scale bar represents 100 μm. **c**. Quantitative measurements of fibrosis (top), and semi-quantitative inflammation scores as assigned by pathologists blinded to the test materials. Mean ± s.d. Significance from unpaired t test.

### 2.4. Extending Lifetime of Osmotic Insulin Pumps

To whether these novel polyacrylamide-based copolymer hydrogel coatings can improve the functional lifetime of insulin pumps, we used implantable osmotic Alzet^®^ pumps. These infusion pumps allow for accurate and continuous delivery of therapeutic formulations, including clinical insulin formulations, allowing us to mimic the continuous basal administration of insulin into the subcutaneous tissue with a traditional infusion set catheter tip used clinically. Flow rate crucially impacts the lifetime and aggregation of proteins on a catheter tip. For humans, typical values of insulin dosing are in the range of 1 U/kg/day^27^. We therefore chose an insulin delivery rate of 0.25 μL/hour (0.025 U/hr insulin) to mimic a similar basal rate. Because the standard tips of the Alzet^®^ pumps are made of metal, we attached a silicone catheter tip to the pump tip comparable to clinical infusion sets. These catheter tips were either left uncoated (bare) or coated with our top-performing copolymer hydrogel D5 (**Fig. 5a**). The pumps were primed for 2 days at 37°C in PBS, per the manufacturer’s instructions, and loaded with insulin lispro (Humalog, Eli Lily) prior to subcutaneous implantation in a rat model of insulin-deficient diabetes (induced with streptozotocin, STZ). Following implantation, serum lispro concentrations were monitored by ELISA throughout the 28-day release period of the particular Alzet^®^ pumps used in this study (**Fig. 5b**). We sought to determine when the catheter tips occluded due to biofouling, resulting in pump failure. Upon occlusion, Alzet^®^ pumps rupture, causing an immediate release of the remaining insulin and resulting in a spike in serum lispro levels^28^ (**Fig. 5d, Fig. 5e, Fig. S5**). In these studies, we defined a cutoff for excessive release of insulin at a serum insulin concentration of 10 mU/L based on literature reports (**Fig. 5d**)^40–42^. While all pumps fitted with bare catheters failed within the 28-day study period, all D5-coated pumps remained functional and maintained consistent serum lispro levels. As the insulin infusion rate was chosen to simulate basal insulin dosing, there was no insulin coverage for carbohydrate consumption and thus blood glucose levels were not expected to reflect management of diabetes despite basal insulin infusion (**Fig. S6, S7**).

**Figure 5.**
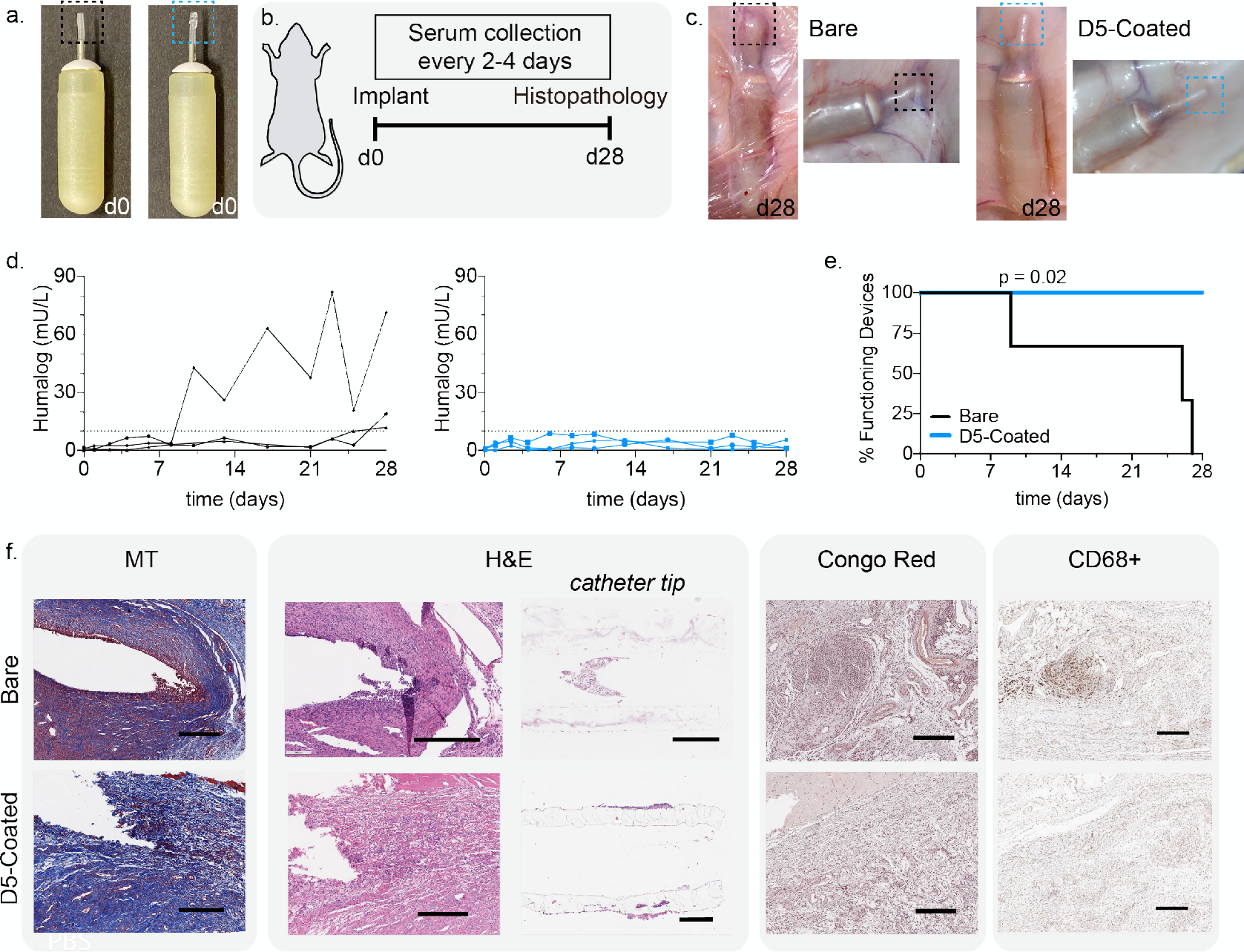
Application of leading hydrogel coating to osmotic insulin pumps. **a**. Osmotic pumps with bare silicone tips (black) and D5 coated tips (blue)**. b**. Pumps are implanted in the dorsal region of diabetic rats, after which pumps and tissues are explanted. **c**. After 28 days, hydrogel coated pumps appear to have noticeably less aggregation. **d**. Blood insulin ELISA **e**. Pump survival curve. Significance assessed with Mantel-Cox test. **f**. Immunostaining and histopathology of excised tissue and catheter tips. Scale bar represents 500 μm.

### 2.5. Biocompatibility of Osmotic Insulin Pumps

At the end of the 28-day study, implanted pumps were retrieved and physically evaluated (**Fig. 5c**). While a thick fibrous capsule and aggregation was observed on bare catheter tips, notably less fibrosis was observed for D5 hydrogel-coated catheter tips. Tissue samples surrounding the tips, as well as the catheter tips themselves, were extracted for histological analysis and stained with MT, H&E, Congo Red, and also immunostained for the macrophage cell market CD68 (**Fig. 5f**). MT staining demonstrated fibrous capsule formation around both bare and D5-coated catheter tips, and several populations of lymphocytes (eosinophils, monocytes, macrophages, and foreign body multi-nucleated cells) were observed around bare catheters. Notably, the region surrounding the bare catheter tips exhibited high levels of inflammation and high density of inflammatory cells (high prevalence of dark blue and purple staining). Moreover, fibroblasts in the region around the bare catheter tips had begun to align — as seen from their nuclear arrangement — as they formed the fibrous capsule around the implanted catheter. These characteristics were absent in D5-coated catheter tips.

Similarly, H&E staining indicated a significantly more inflamed immune response in the tissues surrounding the bare catheter tips than D5-coated catheter tips. Surrounding the bare catheter tips, clear spaces indicated vessels from granulation tissues and capillary tissues, and there was robust fibrotic capsule formation. In contrast, surrounding the D5-coated catheter tips, the deposited collagen was found to be less mature (i.e., recently generated) as a fibrous capsule was beginning to form around the implant. These D5-coated catheter tips also exhibited the presence of inflammatory cells (lymphocytes and macrophages), indicating that while the D5 hydrogel coating does not completely evade the FBR, it remarkably improved the biocompatibility of the implanted catheters.

Additionally, the catheter tips themselves were cross-sectioned and stained, revealing the presence of immune cells on the inside of the bare catheter tips. This cellular accumulation inside the catheter tips could potentially be one driver of the occlusion of the pumps. In contrast, cellular aggregation was not observed inside of the D5-coated catheter tip. Further, Congo red staining indicated amyloid deposition, including from insulin. Aggregates were present in the area surrounding the bare catheter tips, but were negligible in the areas surrounding the D5-coated tips. As macrophages dominate the FBR, CD68 staining was used as a general host immune cell marker to identify macrophages around the implant^29^. The presence of CD68 (macrophage) markers was notably reduced with the D5-coated catheter tips when compared with the bare catheter tips, with 24.5% of the tissue surrounding the bare catheter tips staining positive for CD68 compared to only 15.5% of the tissue surrounding the hydrogel-coated catheter tips (**Fig. 5f**). Commensurate with the results discussed above, these studies indicate that coating of devices with select polyacrylamide-based copolymer hydrogels can reduce the FBR to them upon implantation in the body and improve their functional lifetime.

In these experiments, many factors were expected to contribute to the biocompatibility of the implants, including the presence of insulin, therefore Alzet^®^ pumps filled with PBS were also evaluated for comparison (**Fig. S8**). Insulin was found to aggravate the FBR to the implanted pumps, as pumps with either bare or D5-coated catheter tips delivering PBS infusion showed more mild inflammation and fewer immune cells than pumps delivering insulin. Similar to the results with insulin infusion described above, D-5-coated catheter tips exhibited improved biocompatibility over bare catheter tips when infusing PBS.

## 3. Discussion

In this work, we created a library of polyacrylamide-based copolymer hydrogel materials to develop novel coatings to improve the biocompatibility of implanted devices. Initially, we generated discs of each hydrogel formulation of interest, which were implanted in mice to screen their ability to mitigate fibrous capsule formation and reduce inflammation resulting fro the foreign body response. High-throughput combinatorial synthesis approaches have gained significant interest recently as they have proven to be exceptionally useful for the discovery of new biomaterials for a variety of biomedical applications^17, 30, 31^. We found that one polyacrylamide-based copolymer formulation in particular, comprising a 50:50 mix of HEAm and MPAm, exhibited excellent biocompatibility following implantation into the subcutaneous tissue of mice over the course of one month. Moreover, when this hydrogel was applied to the surface of a PDMS implant, a notable reduction in fibrous capsule formation and local inflammation was observed at the interface of the implant, which was quantified by capsule thickness and assessed through pathological scoring by pathologists blinded to the test materials.

We also found that the HEAm-co-MPAm copolymer hydrogel can be used as a coating to extend the functional lifetime of insulin pumps in a rat model of type I diabetes. Indeed, all insulin infusion pumps fitted with bare catheter tips occluded and failed within the 28-day study period, while all pumps fitted with catheters coated with our lead candidate hydrogels completely maintained their function. While pumps have been shown to improve glycemic control in people with diabetes^45^, the requirement that insulin infusions sets be replaced and moved to new locations in the subcutaneous tissue every few days is highly burdensome. The ability to prolong the functional lifetime of insulin infusion sets with a simple coating could dramatically improve the lives of people with diabetes.

Our leading candidate HEAm-co-MPAm copolymer hydrogel coatings notably improved the biocompatibility of the implanted pump systems by mitigating inflammation and the FBR. While the most desirable immune response to implanted materials is poorly understood and may depend on many application-specific parameters (e.g., tissue types at implantation site, the composition and mechanics of the implanted materials, and the infusion of therapeutics protein formulations to name a few), our studies indicate that our leading copolymer hydrogels may provide a facile mechanism for improving the biocompatibility of a wide variety of medical materials. These polyacrylamide-based copolymer hydrogels can be even further tuned to optimize porosity and even nanotopography, which are known to effect engagement with immune cells^12^. Our findings can inform the design of facile surface modifications of implanted biomaterials for applications ranging from tissue-engineering to biosensors, and our lead candidate hydrogel compositions are promising for next-generation materials and implanted device development.

## 3. Conclusion

In summary, by leveraging a combinatorial synthesis approach to generate a library of polyacrylamide-based copolymer hydrogels with excellent biocompatibility marked by reduced FBR and local inflammation. We were able to develop a simple-to-apply coating based on these materials that helps mitigate the FBR and improve tolerability of various implanted materials including insulin infusion set catheters. Through these studies we demonstrated that our leading candidate polyacrylamide-based copolymer hydrogel coatings have the potential to improve device function and lifetime, thereby reducing the burden of disease management for people regularly using implanted devices, such as those with diabetes.

## 4. Materials and Methods

### 4.1. Materials

All materials were purchased from Sigma-Aldrich and used as received, unless specified otherwise.

### 4.2 Hydrogel Synthesis

Pre-polymer formulations containing 20 wt% acrylamide monomers, 1 wt% lithium phenyl-2,4,6-trimethylbenzoylphosphinate (LAP) as photo-initiator, and 1 wt% N,N’-methylenebis(acrylamide) were mixed in distilled water and pipetted between two glass slides separated by a silicone spacer (0.25 mm ± 0.05 mm). Gels were cross-linked in a Luzchem photoreactor system with 8 W bulbs and an intensity of 25-40 W/m^2^ (LZC-4, *hv* = 365 nm, 15 min). Hydrogels were placed in 1x PBS for at least 24 hr before being punched with a 6 mm biopsy punch. All 2-acrylamido-2-methyl-propane sulfonic acid (AMPSAm) formulations were made with slightly basic solution of 2:3 1M NaOH:water. [tris(hydroxymethyl)methyl]acrylamide (tHMAm) formulations were made with 50:50 dimethylformamide:water as well as 100% N—isopropylacrylamide (NiPAm), 75% diethylacrylamide (DEAm), 25% NiPAm, and 25% hydroxymethylacrylamide (HMAm), 75% NiPAm. PMPC hydrogels was prepared as described previously^15, 23^. Briefly, polyzwitterionic hydrogels were prepared with monomeric solutions in 1M NaCl with 4% N,N’-methylenebisacrylamide (MBAm) cross-linker. A stock solution of 15% sodium metabisulfite and 40% ammonium persulfate was added to the prepolymer solution at ~1wt% and polymerization was initiated at 60°C. PEG hydrogels were prepared similarly with a prepolymer solution of 15 wt% PEG diacrylate (Mn 78024-26) and 5 wt% PEG acrylate (Mn 480).

### 4.3. Platelet Adhesion Assay

Platelet assay was performed as described previously^27^. Briefly, 6 mm punches of hydrogels were incubated for 24 hr at 37 °C with 50% fetal bovine serum (FBS). Then, whole rat blood was mixed with acid citrate dextrose (ACD) anticoagulant buffer for the preparation of platelet-rich plasma (PRP). PRP was obtained via centrifugation at 600 g for 10 min at 10°C. Platelets were diluted to 2.5 x 10^6^ platelets/mL in 1x phosphate buffered saline (PBS). 100 μL of the platelet rich plasma was incubated for 1 hour on hydrogel surfaces. Platelets were rinsed with 1x PBS and fixed with 4% paraformaldehyde (PFA). Hydrogel surfaces were imaged with an EVOS XL Core Imaging System microscope (Life Technologies).

### 4.4. Hydrogel Coating of Polydiemthylsilxoane (PDMS) Discs

Sylgard 184 (Dow) was mixed in 10:1 ratio of monomer:crosslinker and cured at 100°C for 45 minutes per manufacturer instructions. Hydroxyl groups were activated on the surface of PDMS after 3 minutes of O_2_/Ar plasma and 3-(trimethoxysilyl)propyl methacrylate pipetted directly on the surface of PDMS for 10 minutes. PDMS discs were rinsed three times with dIH_2_O and dried with N_2_. PDMS disks were soaked in pre-polymer formulation overnight prior to spin coating, which was done at 500 rpm for 30 seconds before polymerization at 365 nm for 15 minutes.

### 4.5. Scanning Electron Microscopy

SEM images were acquired with an FEI Magellan 400 XHR Microscope with a Beam Voltage of 1 kV and 30 μs dwell time. The sample was pressed onto carbon paint and sputter-coated with Au:Pd (60:40) before imaging.

### 4.6. Ethical Approval of Animal Studies

Animal studies were performed in accordance with the guidelines for the care and use of laboratory animals. All protocols were approved by the Stanford Administrative Panel on Laboratory Animal Care (APLAC-32109; APLAC-32873) and were conducted in accordance with National Institutes of Health guidelines.

### 4.7. In vivo Mouse Biocompatibility

6-8 week-old female B57BL/6 mice were purchased from Charles River Laboratories (Wilmington, MA). Subcutaneous implantations (two per mouse, one on each side of the mouse, and at least 2 cm from point of incision) were made on the dorsal region on mice. Longitudinal incisions (< 2 cm) on the dorsal region was made to access the subcutaneous space. Pockets on either side of the incision were made with a blunt spatula for implanting the materials, one on each side. Incisions were sutured and mice monitored for signs of distress. After 28 days following implantation, mice were euthanized by CO_2_ asphyxiation. The hydrogel and surrounding tissues were harvested (n = 3 for each unique sample) and the tissue was fixed for 48 hr in 4% PFA.

### 4.8. Histology

Fixed tissues were embedded in paraffin and processed by Animal Histology Services at Stanford University. 4 μm cross-sections were stained using Masson’s Trichrome (MT) and Hematoxylin and Eosin (H&E) staining using standard methods. Photographs were taken with EVOS FL Cell Imaging System (Thermo Fisher Scientific) at various magnifications. Rat tissues fixed with 10% formalin for 72 hours and stored in 70% ethanol. Tissues were processed by Histowiz (Brooklyn, NY), fixed in paraffin, sectioned into 5 μm slides, and stained for MT, H&E, anti-CD68 (Abcam, ab125212), and Congo Red.

### 4.9. Pathological Ranking

The thickness for the collagen layer (blue) and pseudosynovium layer (red/purple) were analyzed using ImageJ software, taking the median of 10 measurements per sample. While H&E staining provides a qualitative assay to asses fibrotic response, we sought to quantify the extent to which the hydrogels elicited an immune response. Two pathologists blinded to the test materials graded each H&E stained sample (n = 3 for each hydrogel sample), providing a rating between 0 (little to no inflammatory response) to 3 (severe inflammation), with assessment based on the prevalence of various immune cells at the lining between the skin and implant. Rating was multiplied by area of proportion of impacted area.

### 4.10. Streptozotocin-induced Model of Diabetes in Rats

Animal studies were performed in accordance with the guidelines for the care and use of laboratory animals; all protocols were approved by the Stanford Institutional Animal Care and Use Committee. Male Sprague Dawley rats (Charles River) were used for experiments. The protocol used for streptozootocin (STZ) induction adapted from the protocol by Kenneth K. Wu and Youming Huan^28^. Briefly, male Sprague Dawley rats 160-230g (8-10 weeks) were weighed and fasted 6-8 hr prior to treatment with STZ. STZ was diluted to 10 mg/mL in the sodium citrate buffer immediately before injection. STZ solution was injected into the intraperitoneal space at 65 mg/kg into each rat. Rats were provided with water containing 10% sucrose for 24 hr after injection with STZ. Rat blood glucose levels were tested for hyperglycemia daily after the STZ treatment via tail vein blood collection using a handheld Bayer Contour Next glucose monitor (Bayer). Diabetes was defined as having 3 consecutive blood glucose measurements >400 mg/dL in non-fasted rats.

### 4.11. Preparation of Osmotic Pumps

Commercial Humalog (Eli Lilly) formulations composed of glycerol (1.6%), meta-cresol (0.315%), dibasic sodium phosphate (0.188%), and zinc (0.00197%) were purchased and used as received. Alzet osmotic pumps (Durect Corporation, Model 2004) were filled with ~200 μL Humalog per the manufacturer’s instructions. Pumps had a release rate of 0.25 μL/ hr (0.025 U/hr of Humalog). Pre-filled pumps were placed in sterile saline at 37°C for 48 hr prior to animal implants. For all pumps, 8 mm of RenaSil silicone rubber tubing (Braintree Scientific, SIL037) was attached to the tip of the pumps, due to facile modification of silicone. Hydrogel coated silicone tips were prepared by O_2_/Ar plasma for 3 minutes and immersed in solution of (3-(trimethoxysilyl)propyl methacrylate) for 1 hour, followed by incubation in prepolymer solution for 24 hours. Hydrogel prepolymer solution was pipetted on the tip of the pump and cross-linked for 3 minutes in a Luzchem photoreactor system with 8 W bulbs and an intensity of 25-40 W/m^2^ (LZC-4, *hv* = 365 nm).

### 4.12. In Vivo Implantation of Osmotic Pumps

Following diabetes induction, osmotic pumps were implanted into the diabetic rats. Skin incisions were made on the dorsal region of the rat. A blunt spatula was used to open a pocket in the subcutaneous space. Pumps were inserted and incision closed with 4-0 suture. None displayed visible signs of inflammation or infection throughout the study. After 28 days, animals were euthanized by exsanguination and osmotic pumps and tissue samples were collected.

### 4.13. Blood Glucose and Humalog Quantification

One drop of blood was collected at specified time points via tail vein blood collection onto Contour (R) Next One Blood Gluocse Monitoring System. For determining insulin levels, 60 μL of blood was collected at specified time points via tail vein blood collection into serum-gel microtubes (Sarstedt 50-809-211). Tubes were centrifuged for 5 minutes to extract serum and stored at −80°C. Serum insulin concentrations were determined with Northern Lights Mercodia Lispro NL-ELISA (Mercodia AB, 10-1291-01) according to manufacturer’s instructions, using 10 μL of undiluted serum for each time point. Luminescence was measured in a Synergy H1 Microplate Reader (BioTek). Concentrations were calculated from the generated standard curves.

### 4.14. Statistical Analysis

All implants were conducted with n=3 mice or rats. Unless otherwise specified, measurements are reported as mean ± standard deviation and analyzed with Prism Graphpad v9.0.0.

## Data Availability Statement

The data that support the findings of this study are available on request from the corresponding author. The data are not publicly available due to privacy or ethical restrictions.

## Acknowledgements

Funding: This work was funded in part by NIDDK R01 (NIH grant #R01DK119254) and a Pilot and Feasibility funding from the Stanford Diabetes Research Center (NIH grant #P30DK116074) and the Stanford Child Health Research Institute, as well as the American Diabetes Association Grant (1-18-JDF-011) and a Research Starter Grant from the PhRMA Foundation. C.L.M. was supported by the NSERC Postgraduate Scholarship and the Stanford BioX Bowes Fellowship. We thank the Stanford Animal Histology Services and Histowiz for help with processing of histologic specimens. We thank Dr. José Vilches-Moure (Comparative Medicine, Stanford University) for invaluable discussion on histology images.

## Author contributions

D.C., C.L.M. and E.A.A. designed experiments. D.C., C.L.M. and E.A.A. conducted experiments. D.C., C.L.M, S.S.R., M.L.T. and E.A.A. analyzed data. D.C., C.L.M, S.S.R. M.L.T. and E.A.A. wrote the paper.

