## Supplemental Data for "Polyacrylamide-based Hydrogel Coatings Improve Biocompatibility of Implanted Pump Devices"

### Polyacrylamide-based Hydrogel Coating Improves Biocompatibility of Implanted Devices

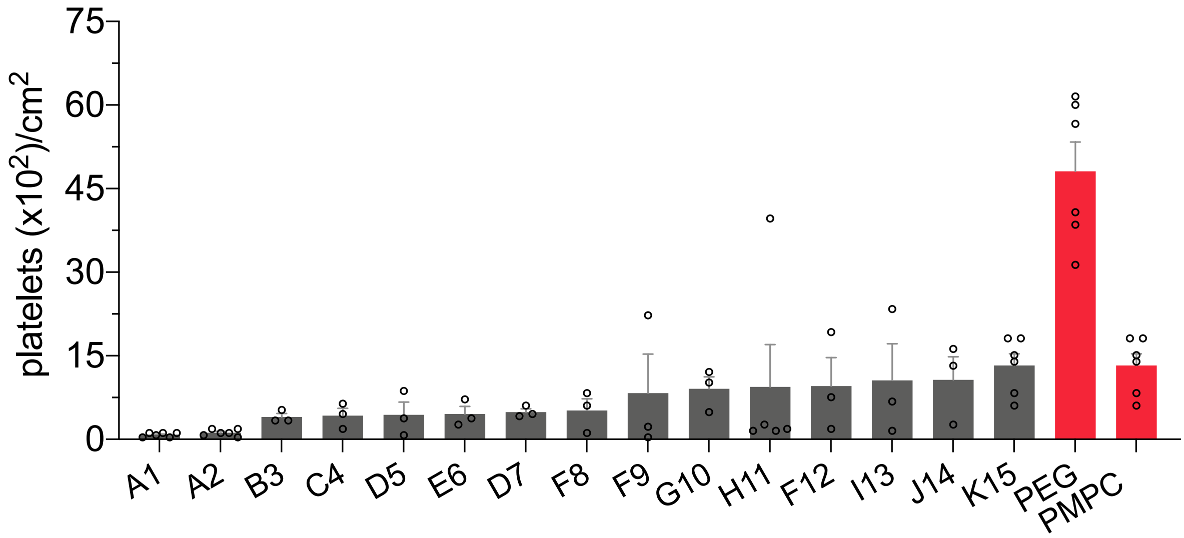


**Fig. S1.** Platelet adhesion counts for top 15 anti-biofouling combinatory polyacrylamide hydrogels and PEG and PMPC hydrogel formulations. Mean ± standard error of platelets on hydrogels are shown with n ≥ 3.

| Ranking on Platelet Adhesion | Monomer 1 | Monomer 2 |
|---|---|---|
| 20 | 15% Am | 5% HMAm |
| 22 | 15% HMAm | 5% MPAm |
| 27 | 20% Am | - |
| 30 | 15% Am | 5% DMAm |
| 46 | 15% NiPAm | 5% Am |
| 49 | 5% ALMP | 15% tHMAm |
| 57 | 15% Am | 5% HEAm |
| 108 | 10% DEAm | 10% tHMAm |
| 151 | 10% ALMP | 10% HMAm |
| 153 | 10% ALMP | 10% DEAm |

**Fig. S2.** Table of formulations from selected mix of anti-blood biofouling hydrogels.

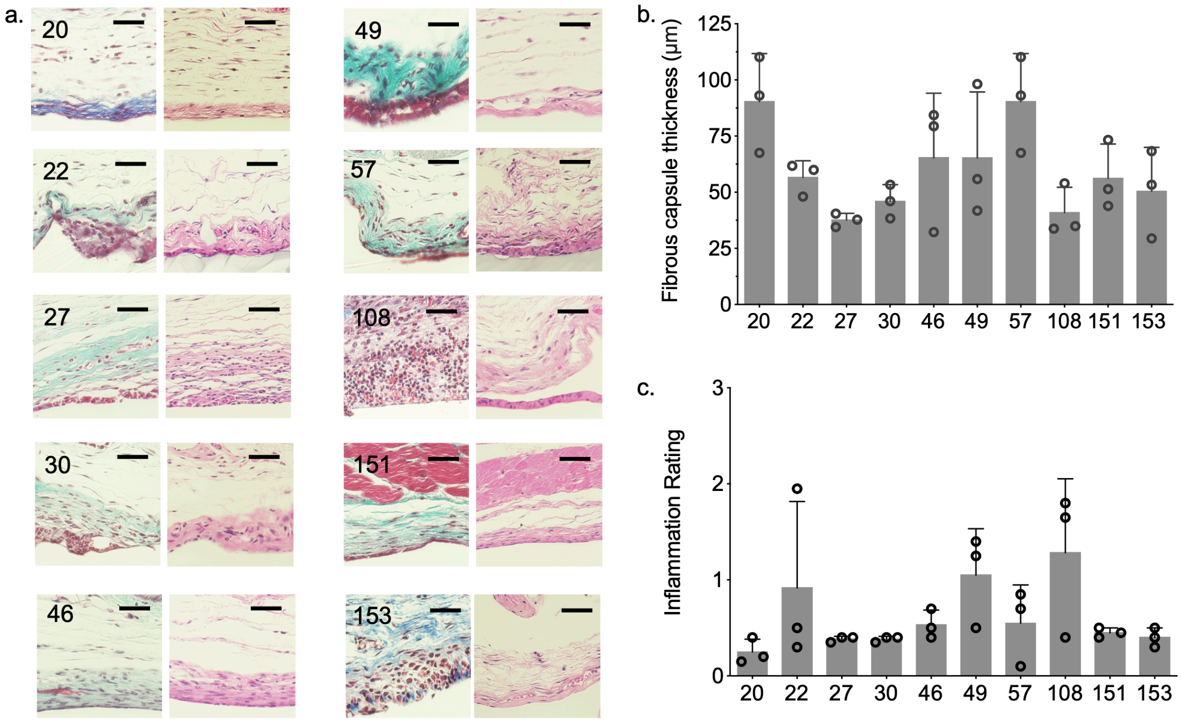


**Fig. S3. a.** Histology images for selected subset of hydrogel library. Number indicates ranking on platelet adhesion test. Scale bar represents 50 μm. **b.** Fibrous capsule thickness of n = 3 samples (mean ± s.d.) where each mean was determined from the median of 10 fibrous capsule thicknesses per image. **c.** Inflammation ratings from blinded pathologists to characterize extent of inflammation.

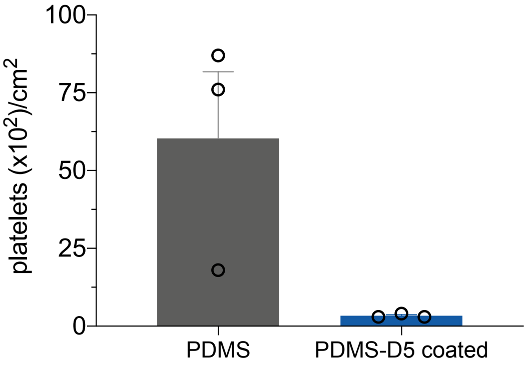


**Fig. S4.** Platelet counts for PDMS and D5-hydrogel coated PDMS. Data shows mean ± standard error of number of platelets on surfaces. Data analyzed with an unpaired t test, p = 0.056, n =3.

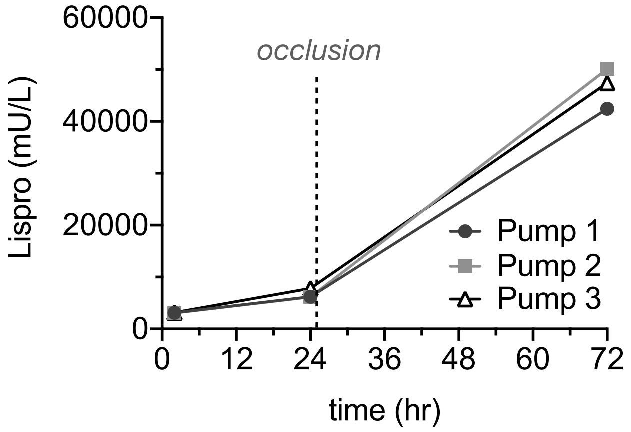


**Fig. S5.** Insulin lispro concentration into PBS from osmotic pumps at 37°C. Aliquots were taken at specified timepoints to determine insulin concentration. Occlusion was induced on the pumps at 24 hr, resulting in a corresponding sharp increase in detected insulin.

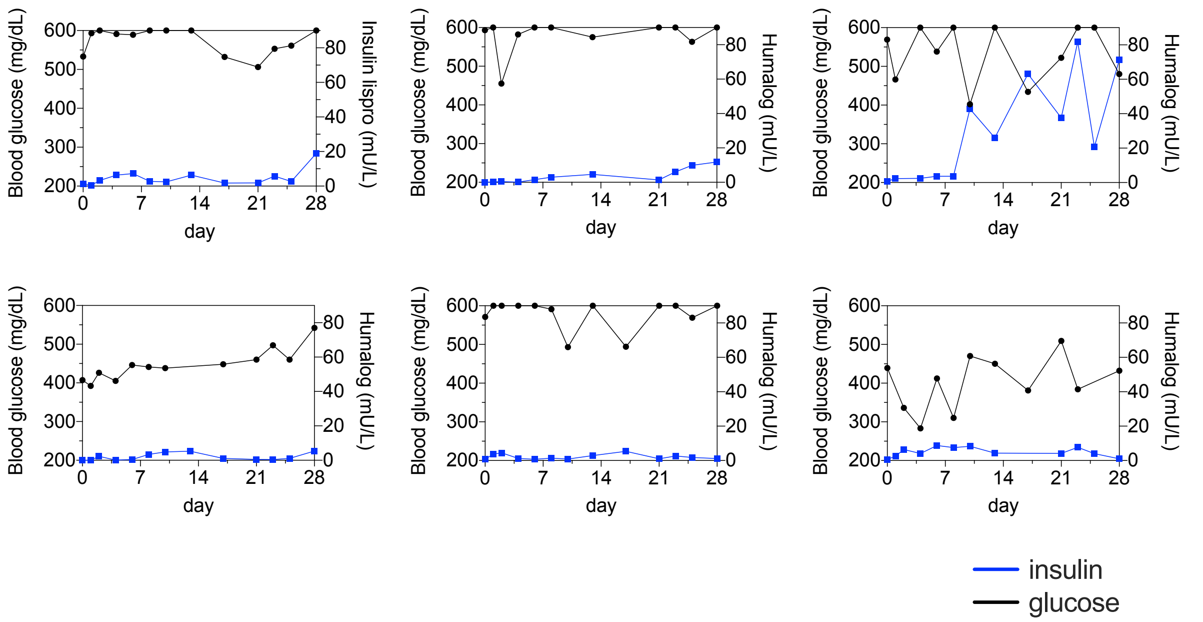


**Fig. S6.** Blood glucose values and humalog concentration from blood serum. Maximum detection limit for blood glucose levels were 600 mg/dL.

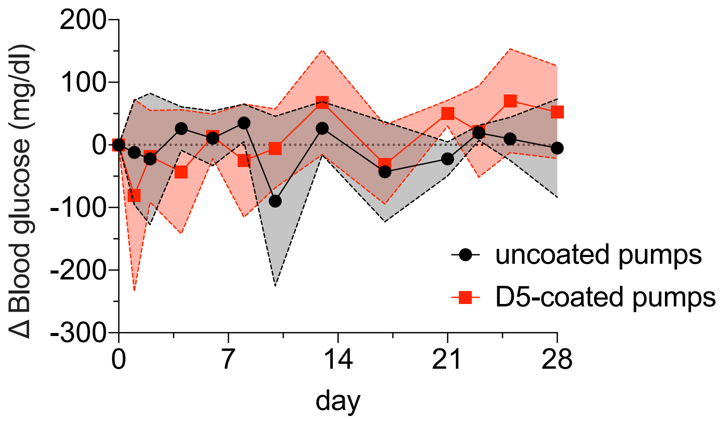


**Fig. S7.** Change in blood glucose levels of rats with uncoated and D5-coated pumps. Data represents mean ± standard deviation, n = 3.

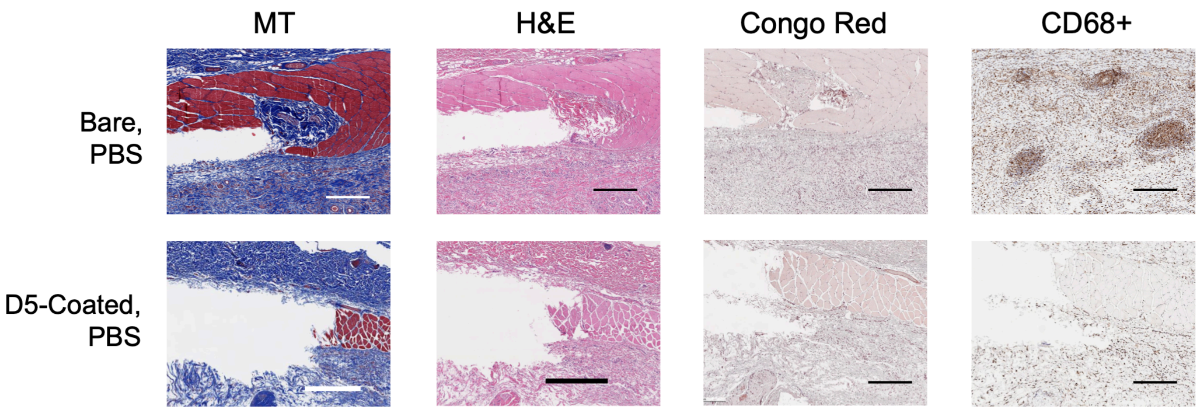


**Fig. S8.** Histology images of tissues surrounding catheter tips (bare and D5-coated) of osmotic pumps infusing PBS. Masson's trichrome (MT), hematoxylin & eosin (H&E), congo red, and CD68+ staining were performed on histology samples, and representatinve images are shown. Scale bar represents 50 μm.
